# Code-Multiplexed Multi-Frequency Impedance Cytometry with a Unified Deep-Unfolding Network

**DOI:** 10.64898/2025.12.13.693849

**Authors:** Wonjun Lee, Sindy K.Y. Tang

## Abstract

Impedance flow cytometry (IFC) is a label-free, single-cell measurement technique that captures biophysical properties beyond traditional biochemical markers. Code-multiplexing allows parallelization of IFC with simple hardware but requires advanced signal processing algorithm to resolve overlaps in signals originating from different channels. Existing methods, however, rely on multiple task-specific networks with template-based linear fitting, which loses accuracy under nonlinear or unstable conditions common in microfluidic experiments. Prior studies have also been restricted to demultiplexing single-frequency impedance measurements. To this end, we develop a unified deep-unfolding network for analyzing code-multiplexed, multi-frequency IFC data. We unfold successive-interference cancellation (SIC) algorithm into a deep-learning network that encodes the structural priors of SIC, and enable a single multitask network to resolve overlaps while sharing information across tasks. To mitigate nonlinear signal stretching and amplification, our network recognizes events by predicting bit-level intensity and duration. For multi-frequency impedance profiling, we apply least-squares fitting to the predicted single-frequency real-impedance trace to map the real and imaginary impedance traces at other frequencies. On the cell-bead mixture evaluation dataset, our pipeline resolves overlaps from singlets to triplets reliably, reconstructs impedance-intensity distributions accurately, and enables multi-frequency impedance profiling. As a demonstration of principle, we use our pipeline to perform label-free quantification of basophil activation from code-multiplexed, multi-frequency IFC measurements. Consistent with previous study, impedance opacity correlates well with activation levels measured by fluorescence flow cytometry. In summary, our study demonstrates the feasibility and utility of a deep-unfolding network that extends code-multiplexed, multi-frequency IFC to label-free single-cell functional assays.

## 1. Introduction

Biological systems exhibit varying degrees of cellular-scale heterogeneity, shaped not only by the surrounding microenvironment but also intrinsic factors such as cell cycle and cell state. Whole blood is a canonical example. It is a complex peripheral system comprising diverse leukocytes, erythrocytes, and platelets spanning orders of magnitude in abundance. Some subsets are rare but have important physiological functions. Basophils, for instance, constitute only ∼1% of circulating leukocytes yet serve as key effector cells in allergic responses (Galli, 2000). Their activation state in response to allergens predicts allergic reactivity and provides clinically actionable information (Castaño et al., 2020; Chinthrajah et al., 2018; Santos et al., 2021). As another example, subsets of tumor cells undergoing epithelial-mesenchymal transition exhibit increased deformability and altered metabolic activities (Hanahan and Weinberg, 2000). Identifying these subsets of cells can reveal metastatic risk and patient outcome, which are often missed by population-averaged measurements. Consequently, extensive efforts have been devoted to quantifying phenotypic heterogeneity at the single-cell level.

Many important phenotypes, such as activation, metabolic shifts, or cytoskeletal reorganization, are fundamentally biophysical rather than purely molecular. Fluorescence labeling detects selected biomolecules but cannot measure a cell’s biophysical properties such as electrical or mechanical state directly. Although traditional fluorescence-based flow cytometry is a mainstay of single-cell analysis, its reliance on labeled markers limits its ability to quantify these properties. In contrast, label-free approaches for biophysical single-cell characterization directly probe the biophysical state of each cell and thus provide strong alternatives to fluorescence-based flow cytometry (Zheng et al., 2013). For example, measurements of cellular electrical properties have been used to quantify biophysical heterogeneity in multiple tumor cell lines (Feng et al., 2022; Honrado et al., 2021), and have enabled applications such as cell counting (Ghassemi et al., 2020), sorting (Li and Ai, 2021), drug screening (McGrath et al., 2020), and cell differentiation monitoring (Haandbæk et al., 2016).

Impedance flow cytometry (IFC), an AC electrical impedance technique derived from the Coulter counter, has emerged as a powerful label-free single-cell characterization method. IFC characterizes single cells electrically by measuring electric-field screening as each cell traverses patterned electrodes in a microchannel. The measured impedance shows frequency-dependent dispersions that can be attributed to subcellular structures, such as the plasma membrane, cytoplasm, and nucleus (Valero et al., 2010). Leveraging this functionality, recent IFC studies have demonstrated the measurements of cell viability (Zhong et al., 2021), drug response (Asphahani et al., 2011), differentiation (Holmes et al., 2009), and activation (Kim et al., 2026).

IFC, however, suffers from an inherent trade-off between sensitivity and throughput of analysis. High sensitivity typically requires a small sensing volume (e.g., by using channel constrictions) to increase the cell’s volume fraction and the impedance contrast between the cell and the surrounding medium. On the other hand, high throughput typically favors wide channels because a larger cross-sectional area lowers hydrodynamic resistance and enables higher volumetric flow at a given pressure, so that more cells can pass per unit time without introducing excessive shear stress or clogging (Jagtiani et al., 2006). Several studies have explored IFC parallelization to overcome the sensitivity-throughput trade-off. The most common approach was to use a multichannel configuration with shared electrodes (Jagtiani et al., 2006). Nevertheless, these devices only parallelized the microfluidic channels for detection while sharing electrodes for all channels. The system suffered from inter-channel crosstalk and failed to identify each signal’s channel of origin, thus falling short of true multiplexing (Zhe et al., 2007). To mitigate crosstalk, some studies dedicated a separate readout electrode to each microfluidic channel (Kim et al., 2013), whereas another study multiplexed the channels by driving each electrode at a distinct frequency (Jagtiani et al., 2011). Although these setups achieved multiplexed operation, their complexity scaled linearly with the number of microfluidic channels because each required a separate set of electrodes and corresponding interfaces.

A code-multiplexed Coulter sensor network addresses the above limitations (Liu et al., 2016). Specifically, patterned electrodes embed a unique code beneath each channel. Particles passing over the electrode array produce bipolar digital signatures akin to those in Code Division Multiple Access (CDMA) networks. By operating up to ten parallel channels and combining with telecommunications-based decoding algorithms, this code-multiplexed approach was effective in increasing the throughput of measurements (Liu et al., 2018).

The code-multiplexing approach simplifies hardware by shifting multiplexing complexity to the software decoding pipeline. Because code-multiplexing uses a single signal transducer, the channel identity and signal intensity should be inferred solely from the measured impedance waveform. When impedance signals from multiple channels overlap, they should be resolved to separate the individual signals from each channel. Realizing the full multiplexing potential requires a robust algorithm for signal separation that is tolerant to noise, drift, and inter-channel interference. Nevertheless, prior algorithms are limited: 1) Both successive-interference cancellation (SIC) algorithm and the reported deep-learning pipelines are based on linear template fitting (Liu et al., 2016; Wang et al., 2019). While this method often suffices in digital telecommunications, it can lose accuracy in nonlinear and/or unstable settings. In microfluidic channels, signals could shift nonlinearly in both time and amplitude due to flow instability and experimental factors. During mechanophenotyping, for example, cells may squeeze through constricted channels of diverse geometries at variable speeds, and may deform during measurements to produce impedance shifts that violate linear superposition (Li et al., 2024). 2) The main task of overlap resolution and signal identification was divided into a region proposal network and a signal classification network, with separate neural networks trained for each task in isolation even though the tasks are closely related (Wang et al., 2021). This siloed design fails to exploit shared information across tasks, leading to poor generalization and inefficiency in scaling and maintenance (Kendall et al., 2018; Vandenhende et al., 2022). 3) The analyses demonstrated thus far remain limited to counting and size estimation at a single frequency near 500 kHz, neglecting the rich biophysical information available from multi-frequency measurements. Taken together, while code multiplexing and neural networks each hold considerable promise, their current integration has yet to realize its full potential.

In this paper, we report a deep-learning pipeline for analyzing code-multiplexed, multi-frequency IFC data. The key innovations of our method are: 1) our deep-learning model recognizes events by predicting bit-level intensity and duration, thereby mitigating nonlinear signal stretching and amplification; 2) the pipeline unfolds the SIC algorithm and harnesses the structural priors of SIC to enable a single multitask network to resolve overlaps while sharing information across tasks; and 3) an auxiliary demultiplexing algorithm enables reconstruction of multi-frequency impedance profiles by propagating predictions from a single frequency to all other frequencies. We demonstrate the effectiveness of our deep-learning pipeline and multi-frequency demultiplexing algorithm using mixed cell-bead datasets obtained from multichannel IFC experiments. As a proof of concept, we apply our pipeline to perform label-free quantification of basophil activation in four parallel stimulation conditions using code-multiplexed IFC measurements. Overall, this work presents a unified, nonlinear, end-to-end network for analyzing code-multiplexed, multi-frequency IFC data that is scalable to high-throughput assay formats.

## 2. Materials and Methods

### 2.1. Device fabrication

We fabricated the IFC device by bonding a polydimethylsiloxane (PDMS, Sylgard 184) microfluidic channel to a glass slide patterned with Ti/Au electrodes, using standard microfabrication and soft lithography techniques. We fabricated electrode-patterned glass slides using a lift-off process. Briefly, we patterned AZ-1512 positive photoresist (MicroChemicals) on a glass wafer using photolithography, and then deposited 5 nm of Ti and 100 nm of Au by electron beam evaporation. After lift-off in acetone with mild sonication, we diced the patterned wafer into individual chips. We fabricated the microfluidic channel in PDMS using soft lithography. We patterned a 15 µm-tall SU-8 master mold on a silicon wafer and then cast and cured PDMS on the mold to form the channel layer. We bonded the PDMS replica to the electrode-patterned glass slide to form the final device following published methods (Tang and Lee, 2010). Briefly, we treated both the PDMS replica and the electrode-patterned glass slide with oxygen plasma for 1 min. We then immersed the PDMS in a 5% (v/v) aqueous solution of (3-aminopropyl)triethoxysilane (APTES, Sigma Aldrich) and, in a separate bath, the glass substrates in a 5% (v/v) aqueous solution of (3-glycidoxypropyl)trimethoxysilane (GPTMS, Sigma Aldrich). After 20 min of incubation at room temperature, we rinsed the substrates with distilled water, dried them with air, brought them into conformal contact using a custom aligner (Fig. S1), and bonded them at 95 °C for 20 min to create the final device (Fig. 1) Fig. S2 details electrode and channel dimensions.

**Figure 1.**
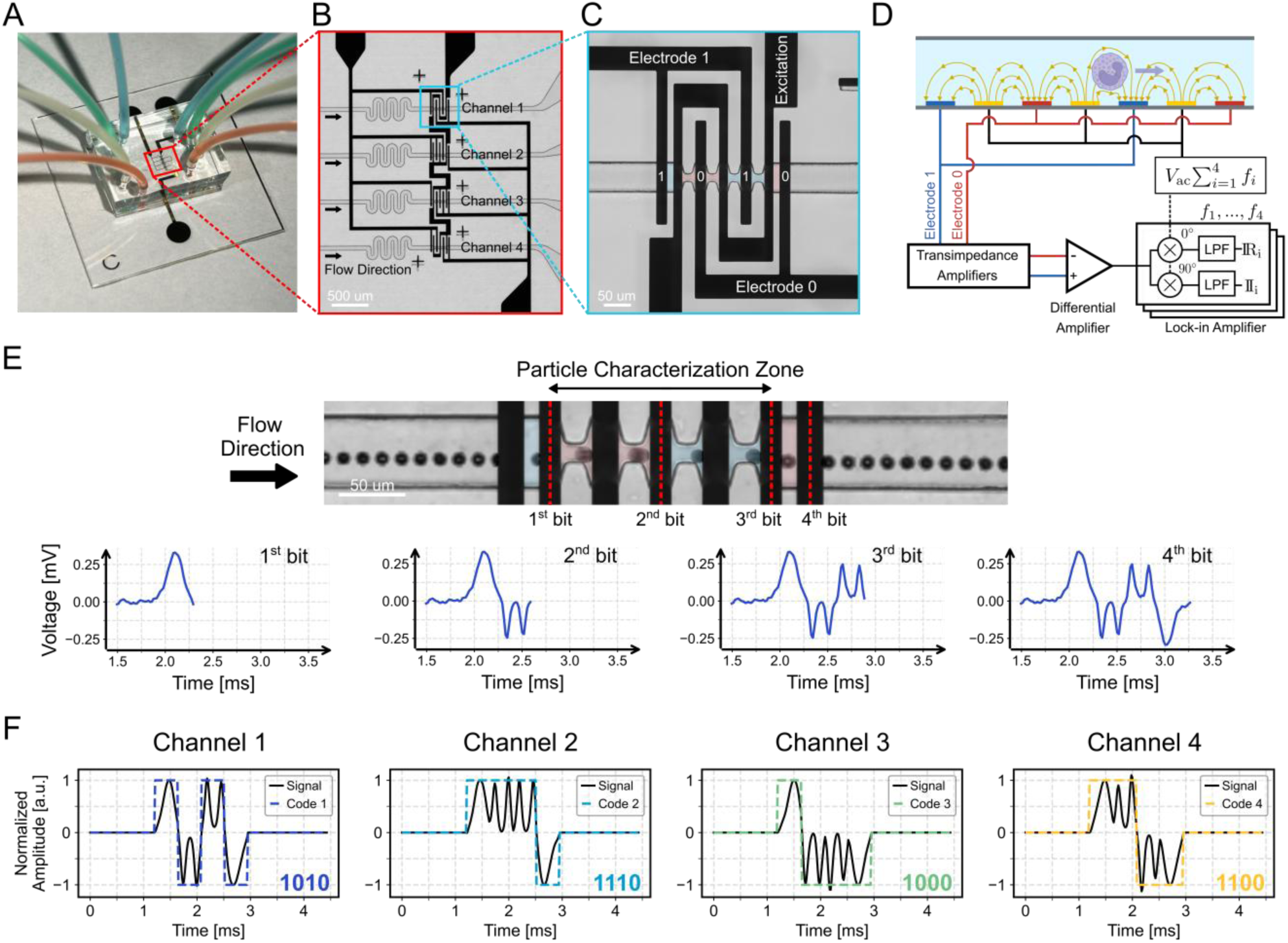
: Impedance flow cytometer (IFC) device design and operating principle. (A) Photograph of the device with each channel filled with a different dye for visualization. (B) Microscopy image of the code-multiplexed IFC device, where gold electrodes were patterned on a glass substrate to form four coded channels with unique electrode patterns. (C) Close-up view of the first channel with the code sequence of 1010. (D) Experimental setup for impedance signal acquisition from the IFC device. *V*_ac_ is the applied AC voltage amplitude, *f*_i_ is the *i*_th_ applied AC frequency, and LPF is a low-pass filter. (E) Time-lapse high-speed images and corresponding impedance signal at 811 kHz of a bead flowing over the electrodes. (F) Representative normalized impedance signals (templates) for each microfluidic channel shown together with the corresponding ideal square-pulse sequences. Templates were generated by averaging training waveforms within each category, normalizing each bit to unit intensity, and zeroing non-signal segments.

### 2.2. Basophil sample preparation

All blood samples were collected by intravenous blood draw into heparinized tubes with informed consent under Stanford University’s Institutional Review Board and in accordance with NIH and FDA guidelines on working with human subjects. We analyzed blood samples from a total of N = 12 donors: four for building the training and evaluation datasets, and the remaining eight for performing label-free basophil activation measurements. Of these 12 samples, the four used for training/evaluation datasets and six of the eight used for label-free basophil activation measurements (samples #1-6, Table S1) were from anonymous, presumed nonallergic donors. Two of the eight samples for label-free basophil activation measurements (samples #7-8, Table S1) were from allergic donors, whose allergic status was confirmed by medical history and skin prick tests (SPT) results.

After blood collection, we stored blood samples at 4°C up to 6 hours before experiments. We isolated basophils from 1 mL of whole blood for each stimulation condition using immunomagnetic negative selection (EasySep™ Direct Human Basophil Isolation Kit) according to the manufacturer’s protocol. We washed the isolated basophils from 4 mL of whole blood in 12 mL of Roswell Park Memorial Institute (RPMI) medium supplemented with 1 mM MgCl₂_(aq)_ and CaCl₂_(aq)_. We then split the cell suspension into four aliquots, and incubated with stimulants for 20 minutes at 37 °C. For N = 4 donor samples used for training/evaluation dataset, the stimulant was anti-IgE (α-IgE) at final concentrations of 0, 10, 100, and 1000 ng mL⁻¹, respectively. For N = 8 donor samples used for label-free basophil activation measurements, the stimuli are listed in Table S1. We quenched stimulation by washing the samples at 4 °C in a low-conductivity buffer. The low-conductivity buffer was prepared by mixing 5 mL 1x PBS with 20 mL of 0.28 M sucrose dissolved in deionized water, then adding 100 µL of 0.5 M EDTA and 417 µL of 30% BSA to reach a final concentration of 2 mM EDTA and 0.5% (w/v) BSA (Invitrogen 15575020; Sigma-Aldrich 126625). The final conductivity of the low-conductivity buffer was measured to be ∼2.9 µS/cm (Thermo Scientific VSTAR50 pH/Conductivity Multiparameter Benchtop Meter). For calibration, we added 8 μm-diameter polystyrene (PS) beads to each sample, achieving a final concentration of about ∼ 10^4^ beads/mL in the aliquot used for IFC measurements. We centrifuged the samples at 400 g for 5 min at 4 °C, discarded the supernatant, and retained approximately 100 μL of basophils (∼ 4×10^3^ cells/mL) and beads for IFC measurements.

### 2.3. Impedance measurements

We injected four aliquots of a donor’s sample, already stimulated with four different stimulants, into four parallel microfluidic channels simultaneously. We injected approximately 100 μL of the cell / bead suspension at a flow rate of 5 μL/min into each microfluidic channel using a syringe pump (Fusion 101, Chemyx Inc.). The event rate was 300-400 events per minute.

We used a lock-in-amplifier (LIA) (HF2LI, Zurich Instruments) and a trans-impedance-amplifier (TIA) (HF2TA, Zurich Instruments) to acquire basophil signals from the IFC device. We applied sinusoidal excitation signals at 101.3 kHz, 811 kHz, 1.13 MHz, and 4.33 MHz to the excitation electrode, with peak amplitudes of 6 V. The resulting basophil signals were collected via the sensing electrodes sandwiching the excitation electrode and amplified through the TIA (resistor R_f_ = 10 kΩ) before being relayed back to the LIA. Within the LIA, the signals passed through a differential amplifier and were synchronized with four demodulators set to the corresponding excitation frequencies. These signals were filtered using a low-pass filter set at 10 kHz, and the final output signal was captured at a rate of 115.1 kHz, which was chosen considering the flow rate at 5 μL/min with a corresponding particle residence time of ∼ 2 ms.

As each cell or particle traversed the measurement region in each microfluidic channel (referred to as “channel” herein), eight characteristic signals were produced, representing the real (𝑥) and imaginary (𝑦) impedance components at four distinct frequencies. We selected the real (𝑥) component at 811 kHz as the triggering signal. When the triggering signal reached a dynamic threshold value, LIA initiated event capture from all 8 signals (real and imaginary voltages at 4 frequencies). We set the dynamic threshold to 120 µV, corresponding to approximately 50% of the first-bit signal peak for beads, 70% for cells, and about eight times the root-mean-square background noise (∼13 µV). The raw data, comprising real (𝑥) and imaginary (𝑦) voltage values at four frequencies, was stored in HDF5 format for post-processing.

### 2.4. High-speed camera image acquisition

For particle imaging, a Phantom V7.3 high-speed camera (frame rate 2000 fps, exposure time 10 μs) connected to the lock-in amplifier was used to capture images of flowing particles. Custom MATLAB code synchronized impedance signal acquisition with camera imaging.

### 2.5. Deep learning for overlap resolution

#### 2.5.1. Data collection and augmentation for deep learning

We collected raw impedance cytometry data from experimental runs for model training and evaluation. Each experimental run was conducted under conditions identical to the label-free basophil activation measurements. We used a new IFC device for each donor. All four channels recorded impedance signals simultaneously from a mixture of PS beads and basophils stimulated with α-IgE at final concentrations of 0, 10, 100, and 1000 ng mL⁻¹, respectively.

Within the training dataset, for each singlet impedance signal measured at 811 kHz, we inspected its waveform and manually labeled the source channel, bit boundary, and the absolute peak intensity of each bit (Fig. 2A). The manually selected bit boundaries and peak positions were used to extract the corresponding bit-intensity and bit-duration labels for 𝑥- and 𝑦- impedance traces across all frequencies. The 811 kHz singlet 𝑥-impedance traces and their labels were used to train and evaluate the deep-learning model, whereas impedance traces acquired at the other frequencies were reserved for assessing the pipeline’s multi-frequency characterization capability. After a second validation pass involving manual re-annotation to verify label accuracy, we retained 2,400 non-overlapping sensor waveforms per channel across four devices (i.e., from four donors). To prevent data leakage, within these 2,400 waveforms, we allocated half (from two donors or devices) to the training of the model and the other half (from the other two donors or devices) to evaluation.

**Figure 2:**
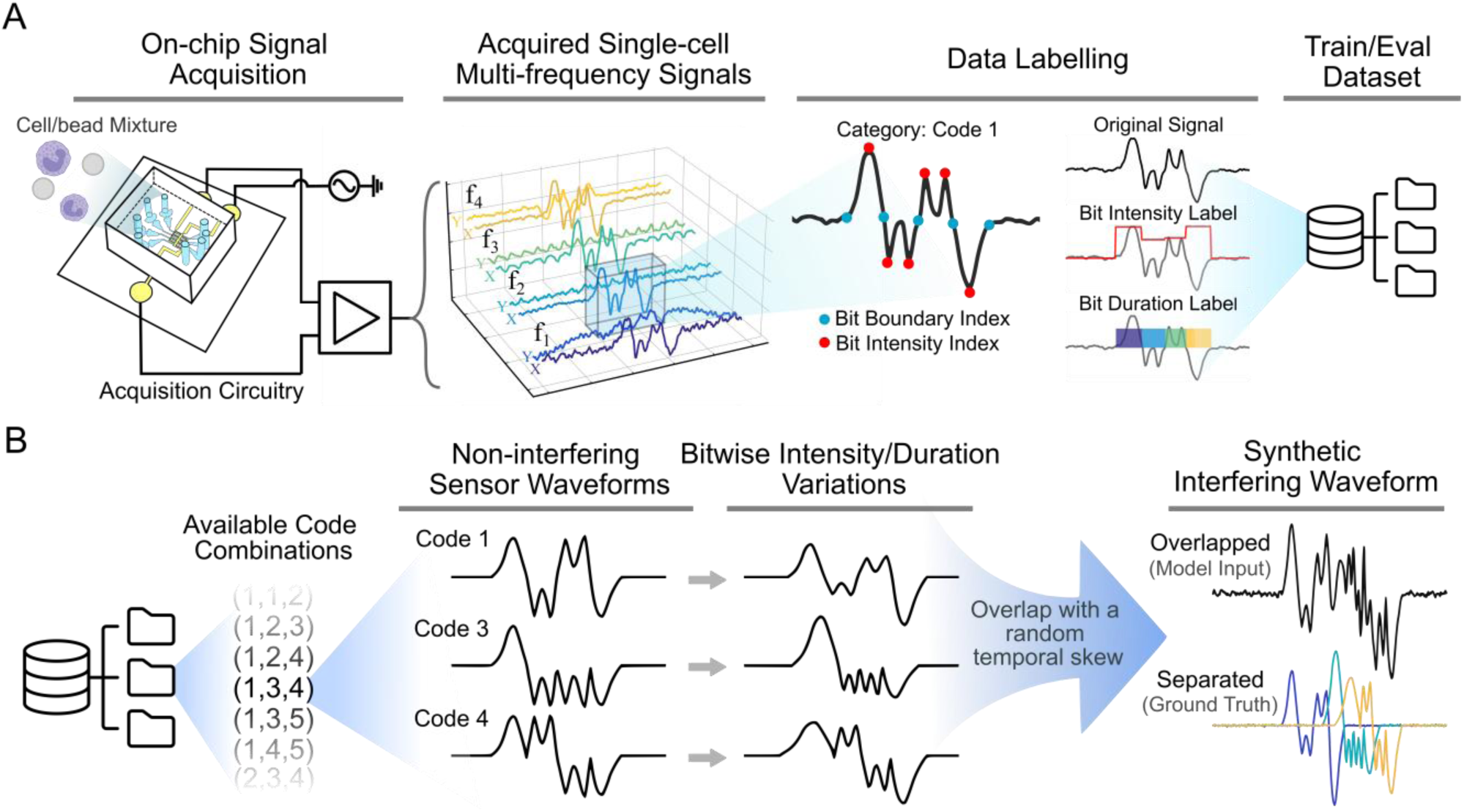
Workflow for the generation of training and evaluation data. (A) Data labeling procedure used to build the training and evaluation datasets. Non-overlapping impedance waveforms recorded from the IFC devices were manually labeled to identify bit-intensity and bit-boundary labels. The resulting labeled traces were stored for use in generating synthetic overlapping waveforms during training and evaluation. (B) Data-augmentation workflow for creating synthetic overlapping waveforms. Singlet waveforms corresponding to each class in the chosen combination were randomly drawn. Each waveform then underwent random amplitude and duration variations for every bit and was temporally offset at random to synthesize an overlapping trace.

Because it is challenging to separate the experimentally acquired overlapping waveforms to identify their constituent signals (ground truth), we generated training and evaluation data by synthetically superimposing non-overlapping waveforms (Fig. 2B). We generated synthetic overlaps up to triplets, which can arise from multiple particles traversing the same electrode in a single channel, or multiple particles traversing different channels at the same time. Assuming four samples at the concentration used in our experiments (∼2×10^4^ particles/mL) were processed in parallel through four channels with Poisson-distributed arrivals, the expected fraction of k-particle events with *k* > 3 is less than 1.62E-4% (*k* = 1: 98.83%, *k* = 2: 1.15%, *k* = 3: 0.02%) (Caselli et al., 2022).

We constructed the database with five signal classes: channels 1, 2, 3, 4 and a no-signal (empty) class. To generate overlapping waveforms, we considered all unique combinations of up to three signals drawn from these five classes (or channels), allowing repetition by enumerating the size-three multiset combinations (with replacement) to yield 35 distinct triplet combinations (e.g., (1,1,1) represents three independent waveforms from channel 1 superimposed; (1,1,2) represents two waveforms from channel 1 and one from channel 2; (1,2,2) represents one waveform from channel 1 and two from channel 2, etc.). For each combination, we randomly selected one waveform from the database for each class in the set and grouped the three waveforms to construct a synthetic overlapping trace. We repeated the procedure until we obtained approximately 100,000 sets that cover a broad range of coincidence scenarios, including singlets, doublets, and triplets.

During training, we applied on-the-fly augmentation by varying the intensity and duration of every bit within each non-overlapping waveform by up to ±25%, and by overlapping the three waveforms with a random temporal skew to synthesize the corresponding overlapping trace. To approximate real measurements, we added experimentally measured, frequency-specific background noise to each synthesized trace. Subsequently, we trained the model to resolve overlaps ranging from singlets to triplets, rather than limiting it to a fixed number of overlaps. This augmentation exposed the network at each epoch to newly simulated IFC device conditions and overlap scenarios, resulting in training on approximately 30 million simulated overlap scenarios, ranging from singlets to triplets, over 300 epochs. Consequently, the network could learn the underlying patterns in heterogeneous overlap scenarios and converge reliably (Fig. S3).

#### 2.5.2. Model architecture

We designed the network architecture to follow the scheme of successive interference cancellation (SIC) (Fig. 3A). We applied a one-dimensional, multitasking UNet (MT-UNet) to the input trace in an iterative manner. At each iteration, the model receives an overlapping trace, identifies the first hidden, non-overlapping trace, subtracts this trace from the input, and passes the residual to the next iteration (Fig. 3B). The iterative UNet comprises a single contracting encoder that is hard-shared by two independent expansive decoder branches. The encoder and decoder layer configurations follow the specifications detailed in the work by Nguyen (Nguyen, 2021). One decoder predicts each bit’s peak intensity through a sigmoid output layer (bit intensity branch, BI), whereas the other predicts bit duration by generating a probability map that assigns every time point to its corresponding bit (bit duration branch, BD). A categorical branch is attached to the bottleneck and connects to multiple layers of the down-sampling path to predict the class (or channel) of the separated trace (trace classification branch, TC). The three outputs—bit intensity, bit duration, and trace class—are combined to reconstruct the hidden trace (Fig. 3C). The intensity-normalized trace template for the predicted class, obtained by averaging all training traces in that category (Fig. 1F), was retrieved and rescaled by an affine transformation with nearest neighbor interpolation to align with the predicted bit positions and durations. The rescaled template trace was then multiplied element-wise by the predicted bit-intensity map, yielding the reconstructed trace for separation. The reconstructed trace is subtracted from the input overlapping trace, and the residual is processed by the same UNet in the next iteration until no overlapping components remain. Under batch-based tensor processing, the unitary prediction time per iteration was 5 ms.

**Figure 3:**
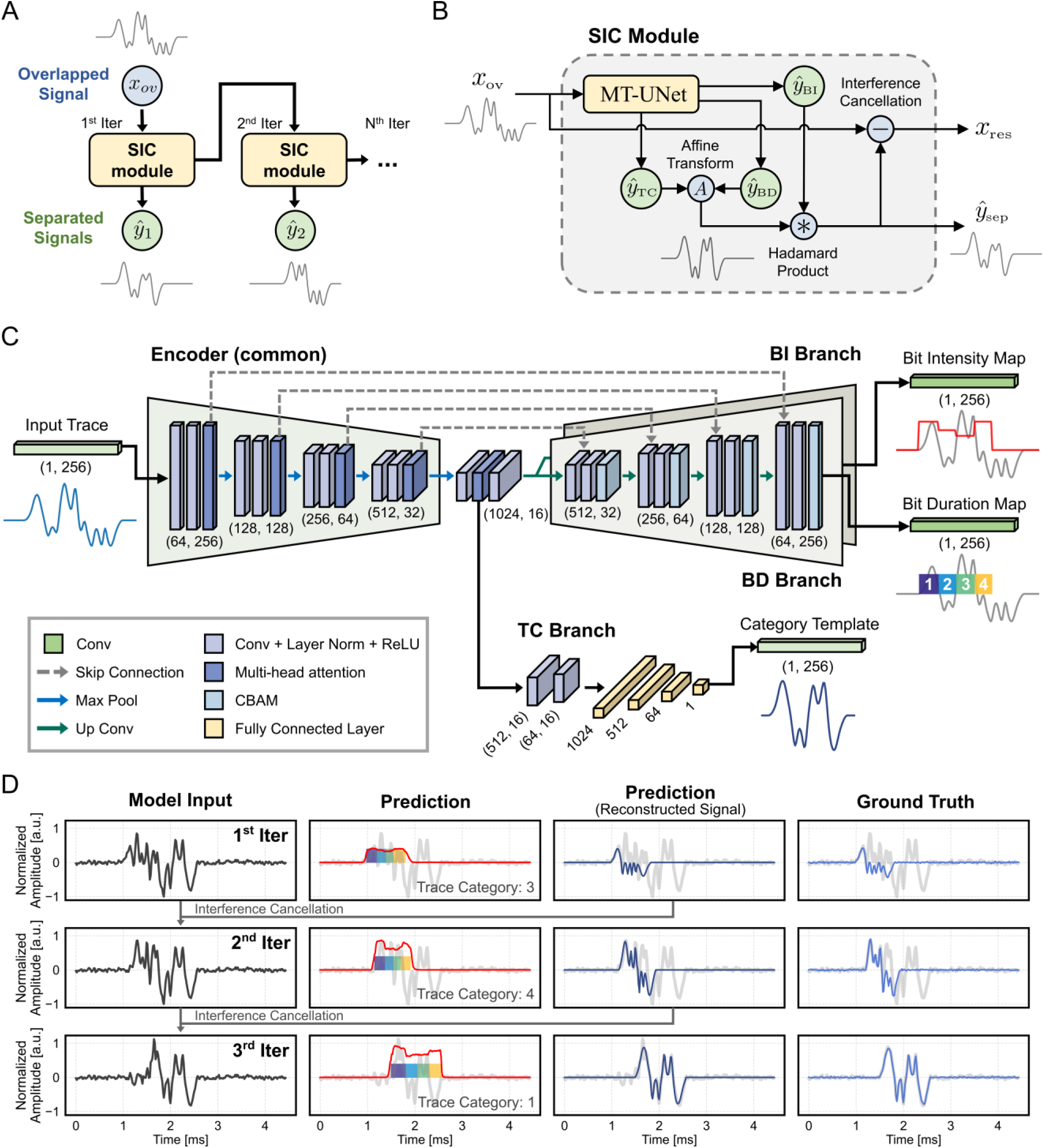
Illustration of the deep-learning pipeline for signal separation. (A) An overlapping input trace was iteratively fed to the successive-interference cancellation (SIC) module, which identified and extracted the first hidden signal and produced an interference-cancelled trace that can be processed again by the same module. We performed up to three SIC iterations. 𝑥_ov_ is the input overlapping trace, and 𝑦̂_𝑘_is the *k*_th_ separated trace. (B) Structure of the SIC module. The input overlapping trace was processed by an MT-UNet that predicted trace class, bit duration, and bit intensity. The hidden trace was reconstructed by retrieving the template corresponding to the predicted trace class and scaling its duration and intensity with the predicted bit-duration and bit-intensity maps. 𝑦_BI_is the predicted bit-intensity (BI) map, 𝑦_BD_is the predicted bit-duration (BD) map, 𝑦_TC_is the category template corresponding to the predicted trace category (TC), 𝑦̂_sep_ is the separated trace, and 𝑥_res_is the overlapping trace after interference cancellation. (C) Architecture of MT-UNet. A single contracting encoder was shared by two expansive decoders, the bit intensity (BI) branch and the bit duration (BD) branch, which predicted the bit-intensity and bit-duration maps of the hidden trace. A categorical trace classification (TC) branch was attached at the bottleneck to predict the class of the hidden trace. Conv is the convolutional layer, ReLU is the rectified linear unit, and CBAM is the convolutional block attention module. (D) Visualization of results obtained with our method for a representative triplet-overlapping trace.

#### 2.5.3. Training scheme

We employ branch-specific loss functions for model training and convergence. The total loss is:

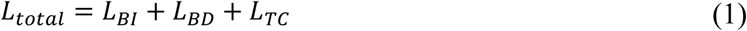

where 𝐿_𝐵𝐼_denotes the loss for BI-branch, 𝐿_𝐵𝐷_denotes the loss for BD-branch, and 𝐿_𝑇𝐶_denotes the loss for TC-branch. Overall, the individual branch losses comprised the following weighted loss functions:

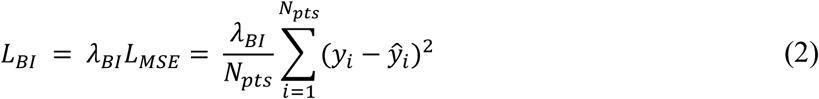

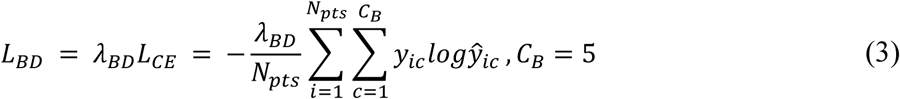

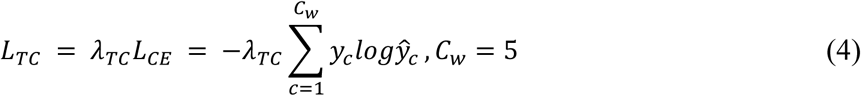

with the contributions of each branch loss controlled by weight hyperparameter 𝜆. In Eq. (2), 𝐿_𝑀𝑆𝐸_denotes the mean squared error for intensity prediction, where 𝑦_𝑖_ is the ground-truth intensity of element *i*, 𝑦̂_𝑖_ is the corresponding predicted value, and 𝑁_𝑝𝑡𝑠_ is the total length of the prediction tensor. In Eq. (3), 𝐿_𝐶𝐸_denotes the cross-entropy loss, which assigns each element to its bit index and classifies the waveform type in Eq. (4). In Eq. (3), 𝑦_𝑖𝑐_is the ground-truth label, 𝑦̂_𝑖𝑐_is the predicted probability that element *i* belongs to class *c*, and 𝐶_𝐵_ is the number of bit categories (first through fourth bit and no bit). In Eq. (4), 𝑦_𝑐_ is the ground-truth label, 𝑦̂_𝑐_ is the predicted probability that trace belongs to class *c*, and 𝐶_𝑊_ is the number of waveform categories (first through fourth channel, and empty signal).

We trained the network on a single NVIDIA RTX 3070 Ti GPU using the stochastic-gradient-descent optimizer with a learning rate of 0.002 and momentum of 0.9. We applied the aforementioned augmentations on-the-fly during training. We set 𝜆_𝐵𝐼_, 𝜆_𝐵𝐷_, 𝑎𝑛𝑑 𝜆_𝑇𝐶_to 10, 1, and 1, respectively, based on empirical tuning so that each branch loss has a comparable magnitude. To promote stable convergence while allowing the network to learn increasingly complex overlaps, we progressively increased the number of iterative passes applied to the overlapping input traces: one pass per sample through epoch 100, two passes through epoch 200, and three passes through epoch 300.

### 2.6. Multi-frequency characterization

For multi-frequency characterization, we first processed the 𝑥-impedance trace measured at 811 kHz with our deep-learning network to identify the underlying non-overlapping trace, which we designated as the pivot trace. We then mapped the extracted non-overlapping pivot trace onto the 𝑥-and 𝑦-impedance traces at the other excitation frequencies, including the 𝑦-impedance trace at 811 kHz, to perform multi-frequency characterization. The identified non-overlapping pivot traces 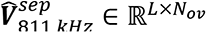 were least-squares fitted to the real and imaginary overlapping traces, 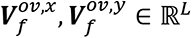, measured at frequency 𝑓 to obtain a frequency-specific linear scaling factor 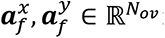

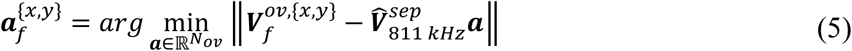

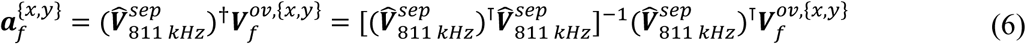

where 𝐿 is the trace length, 𝑁_𝑜𝑣_ is the number of non-overlapping traces identified by the network, and (⋅)^†^ denotes the Moore-Penrose pseudo-inverse operation. Scaling the pivot trace by the factor 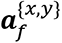 yielded the reconstructed real and imaginary non-overlapping traces at frequency 𝑓:

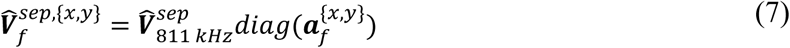

where 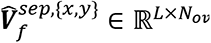.

We evaluated our multi-frequency prediction algorithm on labeled data from two donors used for deep-unfolding network evaluation. To generate synthetic multi-frequency overlapping waveforms, we time-shifted and superimposed impedance traces from multiple events. To keep the overlap condition identical for every frequency and for both the 𝑥- and 𝑦-traces, we applied the same temporal offset to the 𝑥- and 𝑦-traces at all frequencies in each overlapping waveform. Except for the choice of frequencies and impedance components, the overlap fractions and temporal offset conditions matched those used in the deep-unfolding network evaluation.

### 2.7. Signal processing

We post-processed the raw data in four steps: extracting features, identification of basophils and beads, and data normalization. During feature extraction, we analyzed the captured signal from all events. We preconditioned the captured signals by applying baseline correction to the real and imaginary components at every recorded frequency. We processed the 811 kHz 𝑥-impedance trace with our pre-trained deep-learning model to extract non-overlapping traces, and then propagated these traces to the other frequencies using the aforementioned least-squares fitting method. Here, when multiple particles in the same channel arrive with no detectable time gap between them (e.g., a doublet or triplet), they were not further resolved but were identified as a single-particle trace. We excluded these multiplets from analysis using size-based manual gating at 101.3 kHz, as described below.

For each non-overlapping trace, we averaged the predicted intensities of the second and third bits (referred to as “intensity” herein), which incorporated the constricted channels, to obtain a single peak impedance value, thereby obtaining peak impedance values for all frequencies and components. The resulting peak values reflected the electrical properties of all detected particles, including basophils, calibration beads, doublets, debris, and others. We manually gated singlets by examining the 101.3 kHz peak voltage amplitude, which encodes particle size, to exclude doublets and debris (Fig. S4A). Subsequently, the impedance profile at 811 kHz displayed basophils and beads as separate clusters. This separation was driven by their distinct electrical properties (Fig. S4B). We identified the cluster with phase values near zero as polystyrene beads and normalized each basophil peak-impedance value against the corresponding bead value at every recorded frequency:

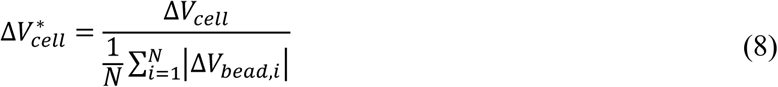

We grouped the resulting normalized impedance profiles by stimulation condition and compared them with fluorescence flow cytometry measurements, which served as the reference for quantifying basophil activation. We assessed the measured impedance profiles using electrical opacity, defined as the ratio of impedance amplitude at a high frequency (*f*_H_) to that at a low frequency (*f*_L_ = 101.3 kHz):

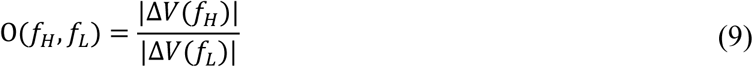

We measured cell electrical diameter as follows:

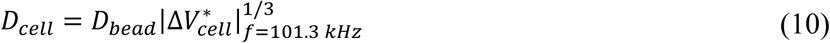

where 𝐷_𝑐𝑒𝑙𝑙_, and 𝐷_𝑏𝑒𝑎𝑑_ are the diameters of cell and polystyrene bead respectively.

### 2.8. Fluorescence flow cytometry

After IFC measurements, we collected basophils from the outlet tubing, stained them, and analyzed them by fluorescence flow cytometry as a benchmark. We stained the collected samples on ice for 30 minutes with antibodies against CD45 (PE), CD63 (APC-Cy7), CD203c (PE/Cy7), and CD123 (APC). After staining, we performed a final wash with the wash buffer (0.5% BSA and 2 mM EDTA in PBS^Ca-/Mg-^), aspirating the supernatant to leave ∼250 μL of the sample in the tubes for fluorescence flow cytometry (Cytek Aurora). Flow cytometry results were analyzed using FlowJo™ v10.8 Software (TreeStar, Ashland, OR). We gated basophils from singlets as CD45^+^/CD123^+^CD203c^+^, and verified through backgating that no stained basophils were excluded from parent gates. We assessed basophil purity after isolation using fluorescence flow cytometry as the fraction of basophils within white blood cells (CD45^+^). The purity was >89% across all experiments with a mean of 96.3% ± 3.4% (see Table S1). We quantified basophil activation as %CD63^+^ basophils using established gating strategies (Pascal et al., 2024). We quantified basophil CD203c activation by subtracting the geometric mean fluorescence intensity (MFI) of the negative control from that of the stimulated sample (Ono et al., 2010):

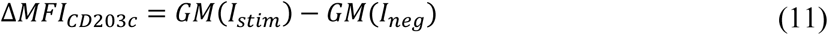

where GM(I) is geometric mean of CD203c fluorescence intensity, 𝐼_𝑠𝑡𝑖𝑚_ and 𝐼_𝑛𝑒𝑔_ are the CD203c fluorescence intensities of the stimulated sample and the negative control, respectively.

### 2.9. Statistical analysis

Cohen’s *d* was calculated to evaluate the effect size among pairwise comparisons (Sullivan and Feinn, 2012).

## 3. Results and Discussions

### 3.1 Device design and operating principle

As a demonstration of principle, we designed the IFC device to incorporate four parallel microfluidic channels with an electrical sensor network embedded beneath each channel (Fig. 1A-D). For the application to a basophil activation test, four parallel channels would be sufficient to test one negative control, one positive control, and two doses of a single allergen or two different allergens. Our electrical sensor network employs the principles of Code Division Multiple Access (CDMA), where the patterned electrodes generate location-specific waveforms to encode each microfluidic channel and multiplex cytometric information into a single electrical output. Consequently, particles flowing through the channels generate impedance signals that reveal both their channel of origin and electrical characteristics.

In code-multiplexed impedance sensing, particles traversing one sensor appear as noise or interference to a decoder tuned to another sensor, thereby lowering the signal-to-noise ratio and ultimately introducing errors in decoding the network’s data. Therefore, we optimized the electrode layout to minimize the bit-error rate introduced by such interference. To reduce the bit error rate, we limited the code length to four bits and omitted flanking excitation electrodes at the outermost bits to minimize the overall sensor length. Previously, by using an improved Gaussian Approximation to estimate the bit-error rate for a correlation-based decoder, it was shown that minimizing sensor length reduced the probability of overlap between particles flowing in different channels and was the most effective in reducing the bit-error rate (Liu et al., 2018). Although our decoder is deep-learning-based rather than correlation-based, both approaches share the same fundamental tasks of interference suppression and signal recovery, so the design guidelines remain applicable.

To increase current density and sensitivity for multi-frequency impedance characterization, we constricted the channel in the regions between the excitation electrode and the sensing electrodes corresponding to the second and third bits (Cottet et al., 2019; Kim et al., 2026). The regions between the excitation electrode and the sensing electrodes corresponding to the first and the last bits were unconstricted to reduce the risk of clogging and were used solely for encoding (Fig. 1C, Fig. S2). Given the design constraints outlined above, we selected binary code sequences that balance the numbers of positive and negative electrode fingers, thereby minimizing baseline current. The chosen sequences are as follows:

Channel 1: 1010; Channel 2: 1110; Channel 3: 1000; Channel 4: 1100.

Fig. 1E shows representative signals and time-lapse images of a bead passing through channel 1 (code 1010). As expected, the particle generated a double-peaked signal as it traversed the second and third bits because here, the excitation electrode surrounded the sensing electrodes. On the other hand, the particle generated a single-peaked signal as it traversed the outermost bits (first and fourth) because the outermost flanking excitation electrode was omitted. Fig. 1F shows representative normalized sensor templates for each microfluidic channel overlaid with the corresponding ideal square-pulse sequences.

While code-multiplexed IFC offers a simple multiplexing interface that requires only a pair of sensing electrodes (0 and 1) and a single excitation electrode for multichannel operation, the extended routing of the excitation electrode can increase its resistance and introduce sensor-to-sensor variation across channels (Fig. S2) (Liu et al., 2018). To assess this potential variation, we injected calibration beads through each channel and compared the measured impedance amplitudes across the four channels. Across all channels from three IFC devices, the standard deviation (SD) of bead 𝑥 - impedance intensities (the mean of the second- and third-bit peak intensities) averaged 15 μV (101.3 kHz: 13 μV; 811 kHz: 16 μV; 1.13 MHz: 16 μV; 4.33 MHz: 16 μV), and the SD of 𝑦-impedance intensities averaged 4 μV (101.3 kHz: 7 μV; 811 kHz: 1 μV; 1.13 MHz: 2 μV; 4.33 MHz: 7 μV). These SD values were comparable to or smaller than the background noise (101.3 kHz: 31 μV; 811 kHz: 13 μV; 1.13 MHz: 23 μV; 4.33 MHz: 14 μV) (Fig. S5). These results confirmed that the measured impedance signals were sufficiently uniform across channels. Furthermore, by applying channel-specific bead normalization, we suppressed residual sensor-to-sensor variation during signal processing (Fig. S6) (Apichitsopa et al., 2018). To further validate that the impedance profiles reflect the electrical properties of the cells rather than device-induced bias, we injected unstimulated basophils (samples #5 and #6, Table S1), through all channels and measured the impedance signal. The mean coefficient of variation (CV) in opacity across the four channels was 2.8% (𝑂(𝑓_𝐻_ = 811 𝑘𝐻𝑧, 𝑓_𝐿_ = 101.3 𝑘𝐻𝑧) : 3.35%; 𝑂(𝑓_𝐻_ = 1.13 𝑀𝐻𝑧, 𝑓_𝐿_ = 101.3 𝑘𝐻𝑧) : 1.5%; 𝑂(𝑓_𝐻_ = 4.33 𝑀𝐻𝑧, 𝑓_𝐿_ = 101.3 𝑘𝐻𝑧): 3.54%) (Fig. S7).

### 3.2 Overlap resolution with deep-unfolding network

Fig. 2 shows the data collection and augmentation procedures for model training, and Fig. 3 shows the network architecture (see details in section 2.5.2). The network architecture unfolds the SIC algorithm as an iterative application of a single learnable SIC module, whose core is an MT-UNet backbone that performs overlap resolution and signal reconstruction. The SIC module takes an input overlapping trace, identifies the first hidden non-overlapping trace, performs interference cancellation, and passes the residual to the next iteration. When the input trace no longer contains any non-overlapping trace, the MT-UNet within the SIC module classifies the hidden-trace category as empty and terminates overlap resolution. This approach avoids the need for separate networks for each overlap scenario (Fig. 3A-B). The MT-UNet identifies individual traces through output branches that predict each trace’s bit-wise intensity (BI), bit-wise duration (BD), and trace category (TC) (Fig. 3C). The shared encoder among branches enables cross-task information sharing. By predicting intensity and duration bit-wise, the model directly addresses nonlinearity commonly present in measured impedance signals in microfluidic experiments and improves the accuracy of signal reconstruction. This feature distinguishes our model from prior methods that only linearly stretch or amplify a single template, ignoring bit-wise signal variations. Fig. 3D shows representative predictions for a triplet-overlapped trace.

Fig. 4 presents the evaluation results for our deep unfolding network. We evaluated the model on labeled data from two donors not used for training. The evaluation dataset comprised synthetic waveforms generated by overlapping 1,200 labeled traces for each code into singlet, doublet, and triplet combinations. We created these overlaps without altering bit-level intensity or duration. We assessed whether the intensities of reconstructed signals were accurate and whether the TC branch assigned each signal to its class correctly. The results show that the TC branch achieved mean classification accuracies of 99.85 % for singlets, 96.93 % for doublets, and 88.83 % for triplets. Among correctly classified signals, 811 kHz 𝑥-impedance intensity exhibited mean error rates of 5.41 % for singlets, 8.70 % for doublets, and 12.68 % for triplets. These results confirm that our pipeline handled overlap scenarios ranging from singlets to triplets flexibly and reconstructed the hidden traces accurately. As expected, the classification and reconstruction accuracies declined from singlets to triplets because increasing particle overlap increased the likelihood that the interference would dominate the target signal. Interference dominates when the target signal’s autocorrelation drops below its cross-correlation with interferers, a condition formalized in digital telecommunications (Holtzmann, 1992).

**Figure 4:**
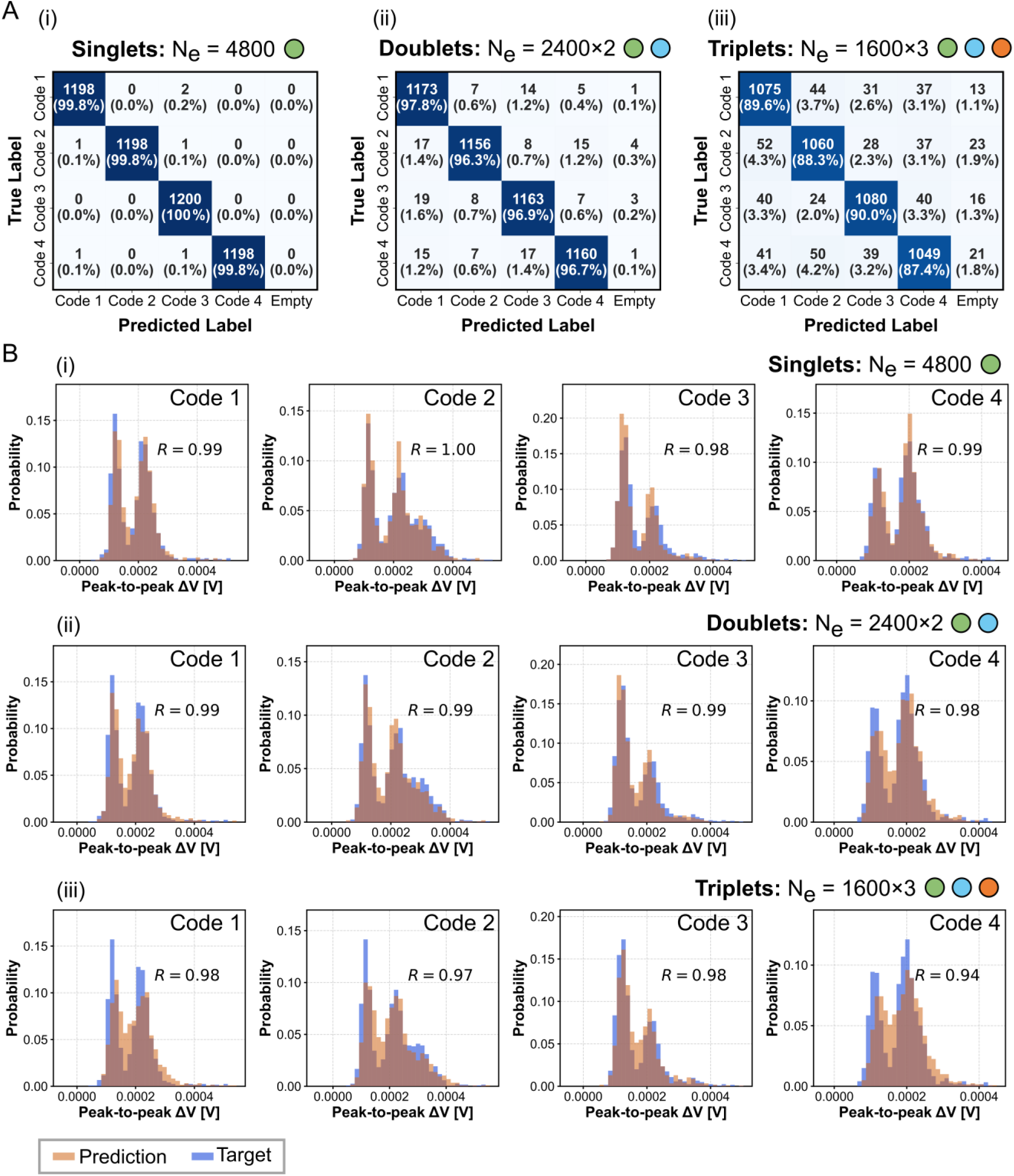
Overlap-resolution results for the in-phase (𝑥) 811 kHz traces obtained with the deep-learning network. (A) Confusion matrix for trace classes (codes 1 to 4) predicted by the trace classification (TC) branch. (B) Histograms comparing the target and predicted intensity distributions of reconstructed traces for each code, obtained from singlet, doublet, and triplet overlaps. Prediction results for singlet, doublet, and triplet cases are shown in panels (i)-(iii). N_e_ is the number of processed non-overlapping traces. Pearson correlation coefficients (R) between the predicted and target distributions are reported within the plots.

### 3.3 Multi-frequency impedance profile prediction

For multi-frequency characterization of individual particles in code-multiplexed IFC, we processed the 811 kHz 𝑥-impedance trace with the deep unfolding network and used it to infer impedance traces at other frequencies. The differential impedance signal measured by IFC can be expressed theoretically by the following equation, derived from the Maxwell mixture model:

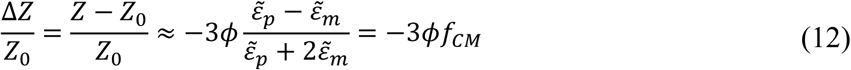

where 𝜙 is the particle electrical volume fraction, 𝑓_𝐶𝑀_ is the Clausius-Mossotti factor, and 𝜀̃_𝑚_, and 𝜀̃_𝑝_denotes the complex permittivities of the buffer medium and particle, respectively (Valero et al., 2010). In Eq. (12), 𝜀̃_𝑚_and 𝜀̃_𝑝_are frequency-dependent material properties and are constant with respect to particle location. Thus, 𝜙 is the primary factor that shapes the impedance waveform as a particle traverses the electrode. Waveforms measured at different frequencies share the same shape and differ only in intensity as determined by the frequency-dependent Clausius-Mossotti factor 𝑓_𝐶𝑀_.

Traces measured at different frequencies should therefore be linearly related, with each component derivable from the 811 kHz prediction by a frequency-specific scaling factor (Eq. (7)). Accordingly, the individual non-overlapping 𝑥-impedance traces at 811 kHz, generated by our deep unfolding network, can serve as templates for inferring signals at other frequencies. We used least-squares fitting to estimate linear coefficients that best map the non-overlapping 𝑥-impedance traces at 811 kHz to the measured overlapping traces at other frequencies (see details in section 2.6). We then applied these fitted coefficients to scale each non-overlapping trace at 811 kHz in intensity and obtain its multi-frequency profile.

Fig. 5 plots the multi-frequency impedance profiles inferred by our method on the evaluation dataset, and Table 1 summarizes the corresponding results (see Supplementary Note S1 for definitions of the evaluation metrics). We computed the intensity error rate by comparing predicted intensities with ground-truth intensities, with the analysis restricted to signals with ground-truth values exceeding three times the background noise level. As shown in Table 1, for frequency components that exhibit a clear shape linearity (𝑅^2^ ≥ 0.94 ) with the 811-kHz 𝑥 -impedance trace (i.e., 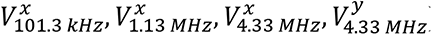), our method yielded intensity error rates within 2% of the deep-unfolding network’s 811-kHz 𝑥 -intensity predictions across codes and overlap scenarios. Frequency components that show reduced linearity (𝑅^2^ ≤ 0.9) with the 811 kHz 𝑥-impedance trace 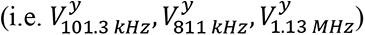 exhibited increased intensity error rates. We attributed the reduced linearities at these frequencies to the low SNR (𝑆𝑁𝑅 < 4) from an attenuated out-of-phase current (i.e., the 𝑦-impedance traces), caused by the particles’ (cells and beads) weak capacitive property at these frequencies (Fig. S8). Nevertheless, with all events included irrespective of SNR, the reconstructed and ground-truth 2D x–y impedance intensity distributions (Fig. 5) at each frequency were well-correlated (Pearson correlation coefficient R ≈ 0.89) across codes and overlap scenarios.

**Figure 5:**
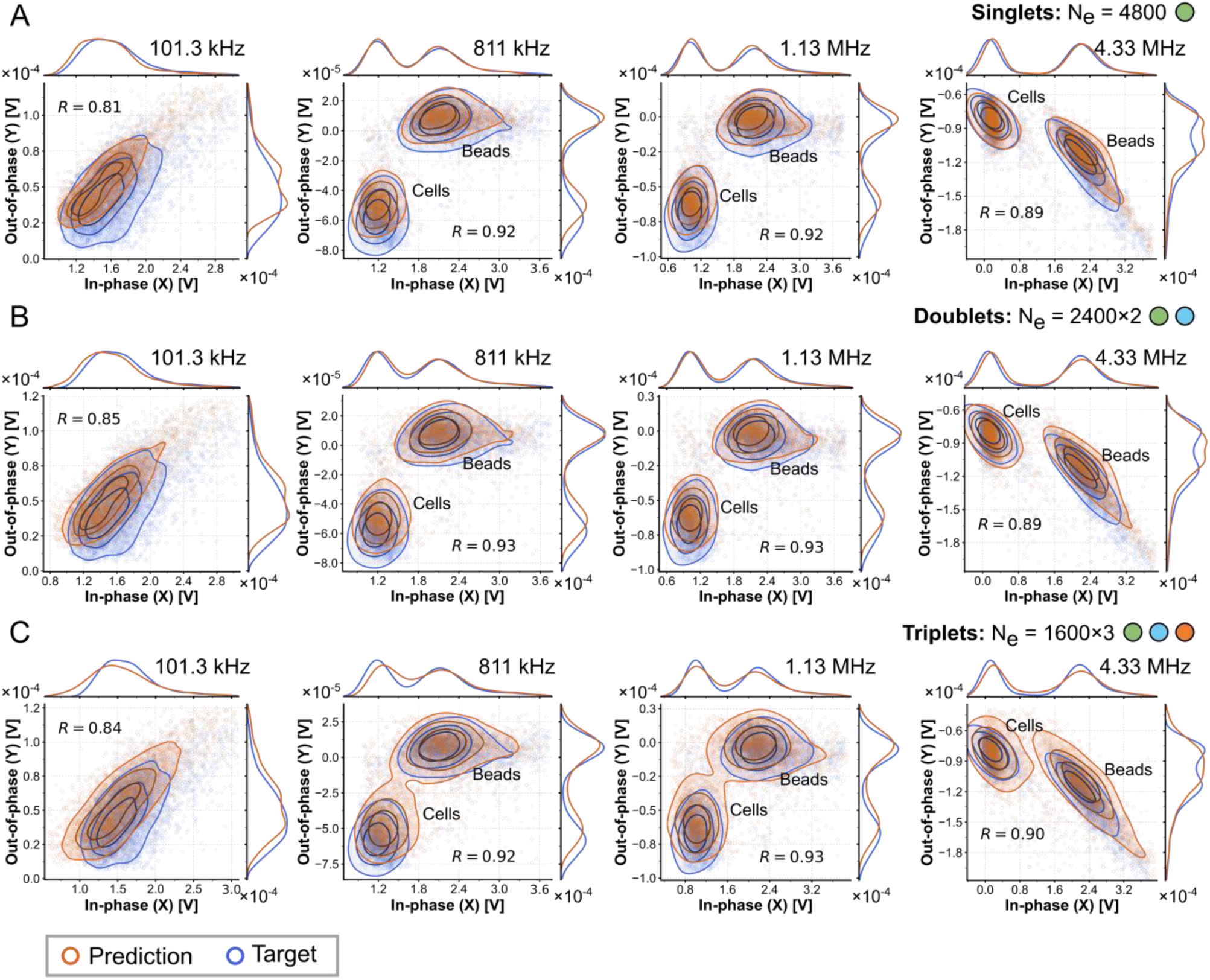
Comparison of the target and predicted intensity distributions of the in-phase (𝑥) and out-of-phase (𝑦) impedance traces at 101.3 kHz, 811 kHz, 1.13 MHz, and 4.33 MHz for singlet, doublet, and triplet overlaps. The in-phase 811 kHz traces predicted by the deep-learning network were used to reconstruct the impedance profiles at the other frequencies. Prediction results for (A) singlet, (B) doublet, and (C) triplet cases are arranged in separate rows. The contour lines denote the 25^th^, 50^th^, 75^th^ percentiles of each population group, from the darkest to lightest shade. Pearson correlation coefficients (R) between the predicted and target distributions are reported within the plots.

**Table 1.** Evaluation results for multi-frequency impedance-intensity prediction. 𝑅^2^was computed for each frequency component with respect to the 811 kHz 𝑥-impedance trace. SNR denotes signal-to-noise ratio. All values were averaged across codes (see Supplementary Note S1 for definitions of the evaluation metrics). Evaluation at the 811 kHz 𝑥-trace (blue) reflects the performance of the deep-unfolding network in resolving overlaps, whereas the evaluation at other frequencies reflects the performance of the least-squares mapping from the 811 kHz 𝑥-trace.

| | $V_{101.3 \text{ kHz}}^x$ | $V_{101.3 \text{ kHz}}^y$ | $V_{811 \text{ kHz}}^x$ | $V_{811 \text{ kHz}}^y$ | $V_{1.13 \text{ MHz}}^x$ | $V_{1.13 \text{ MHz}}^y$ | $V_{4.33 \text{ MHz}}^x$ | $V_{4.33 \text{ MHz}}^y$ |
| --- | --- | --- | --- | --- | --- | --- | --- | --- |
| **Average SNR** | 6.89 | 3.73 | 13.98 | 3.99 | 7.77 | 3.44 | 16.05 | 8.18 |
| **Average $R^2$** | 0.96 | 0.9 | - | 0.86 | 0.94 | 0.79 | 0.98 | 0.95 |
| **Average Error [%]**<br>**(Singlet)** | 7.91 | 12.88 | 5.41 | 16.1 | 7.03 | 17.16 | 6.85 | 7.97 |
| **Average Error [%]**<br>**(Doublet)** | 10.02 | 13.47 | 8.7 | 17.7 | 10.3 | 18.14 | 9.7 | 10.82 |
| **Average Error [%]**<br>**(Triplet)** | 13.41 | 15.84 | 12.68 | 20.96 | 13.93 | 22.47 | 12.5 | 13.79 |

This result indicates that our method recovered the population-level distribution of impedance intensities accurately. Taken together, this agreement indicates that our pipeline is reliable in using the network’s prediction at a single frequency (811 kHz) to infer the corresponding signals at other frequencies to enable multi-frequency impedance profiling for each particle.

### 3.4 Label-free measurement of basophil activation with code-multiplexed IFC

As a proof of concept, we applied our pipeline to quantify basophil activation in a label-free manner using code-multiplexed IFC measurements (Fig. 6). We analyzed blood samples from N = 6 donors: four anonymous, presumed nonallergic donors and two allergic donors (samples #1-#4 and #7-#8, Table S1). All donors were held out from model training and evaluation. Each donor’s sample was split into four aliquots and stimulated at varying concentrations of anti-IgE or food allergens (see Table S1). The four aliquots, together with calibration beads, were then injected in parallel into the microfluidic channels. Impedance profiles were recorded as the suspension flowed through the channels. Samples from each channel were collected and analyzed by fluorescence flow cytometry to measure the expression level of activation markers CD63 and CD203c. Following allergen stimulation, CD63 and/or CD203c upregulation relative to the negative control indicates basophil reactivity and correlates strongly with clinical allergic response to that allergen (Santos et al., 2021). These results were compared with their respective impedance profiles. Because individual events measured by IFC could not be directly matched to those measured by fluorescence flow cytometry, we performed comparisons at the population level following our published approach (Kim et al., 2026).

**Figure 6:**
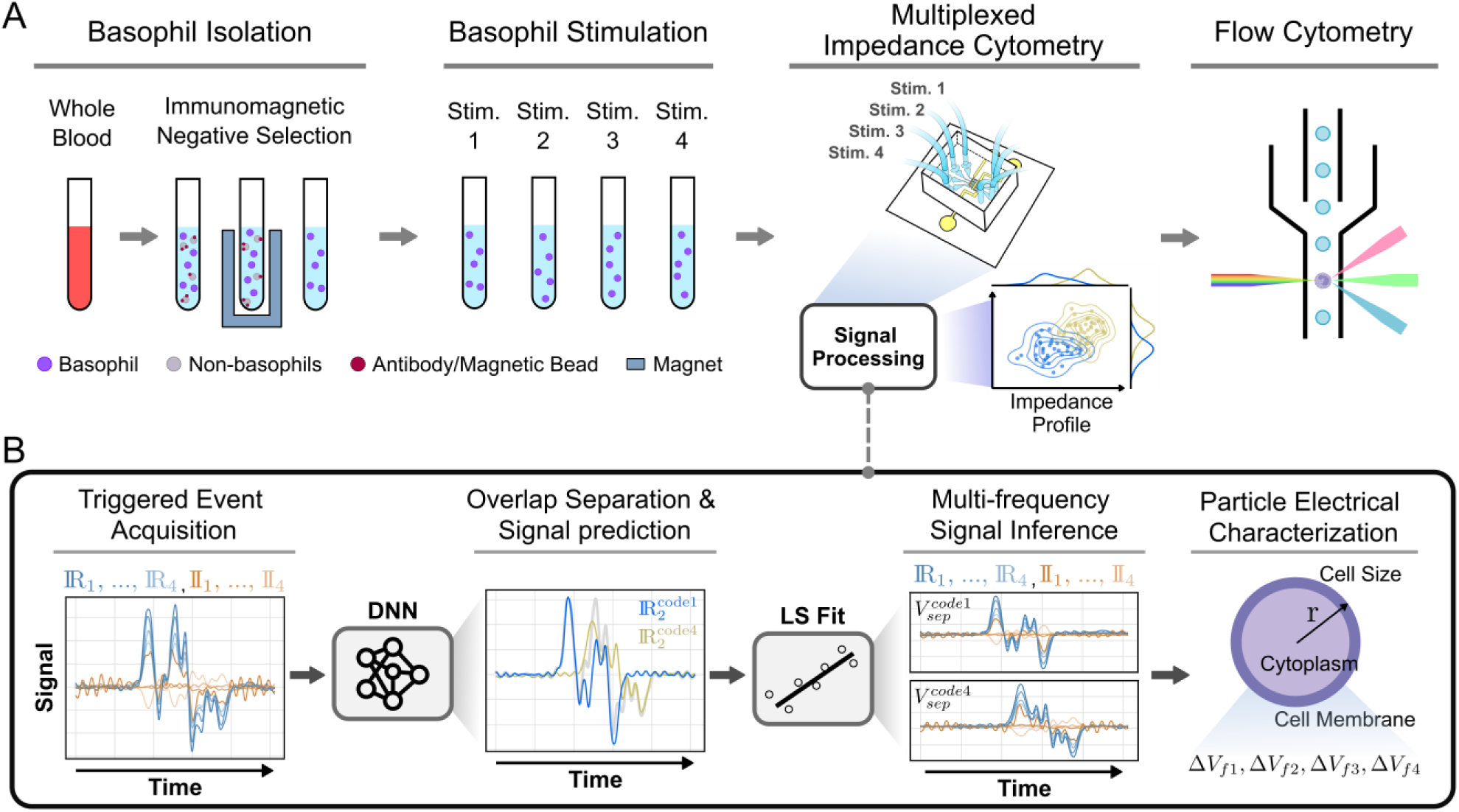
Scheme for experimental label-free measurement of basophil activation using a code-multiplexed IFC device. (A) Basophils isolated from whole blood by immunomagnetic negative selection were stimulated with different stimulants and analyzed in the IFC device, with each microfluidic channel dedicated to a specific stimulation condition. Samples collected from the outlets were examined by fluorescence flow cytometry. (B) Captured multi-frequency impedance signals were processed by a pre-trained deep-unfolding network that used the 811 kHz in-phase (𝑥) trace to identify each signal and to separate any overlaps. Non-overlapping impedance traces at the other frequencies were then inferred from the predicted 811 kHz trace by least-squares (“LS”) fitting. Average peak amplitudes of the second and third bits in each non-overlapping trace at each frequency were used to characterize the measured particles electrically. ℝ_𝑖_ and 𝕀_𝑖_are the real and imaginary components of the measured impedance at frequency *f*_i_, and DNN is the deep-neural network.

Fig. 7A-B shows basophil opacity and diameter from three representative donors. For the nonallergic donor (SBC255), we observed decreasing basophil opacity with increasing α-IgE dose (also see Fig. S9). This result is consistent with our prior observation that activation-induced changes in membrane capacitance reduce opacity. For the two allergic samples (PR0783, PR0784), we stimulated the blood with two food allergens, one that the donor was allergic to and one they were not. As shown in Fig. 7A-B, basophil opacity decreased only with the allergen to which the donor was allergic. Within each donor, stimulation conditions that produced larger differences in CD63 and CD203c activation levels also showed correspondingly larger differences in opacity, as quantified by Cohen’s d effect size (Fig. 7B). Fig. 7C-D compiles the changes in basophil opacity (*f*_H_ = 811 kHz, 1.13 MHz, and 4.33 MHz respectively, see Eq. (9)) from all N = 6 donors in response to different stimulants versus the corresponding activation levels measured by fluorescence flow cytometry. We defined the change in opacity (ΔO) as the difference between the mean opacity of basophils in the stimulated conditions (with anti-IgE- or allergen) and that in the negative control (with no stimulants). Across all frequencies tested, an increase in activation level correlated with a general decrease in basophil opacity, consistent with our previous findings (Kim et al., 2026).

**Figure 7:**
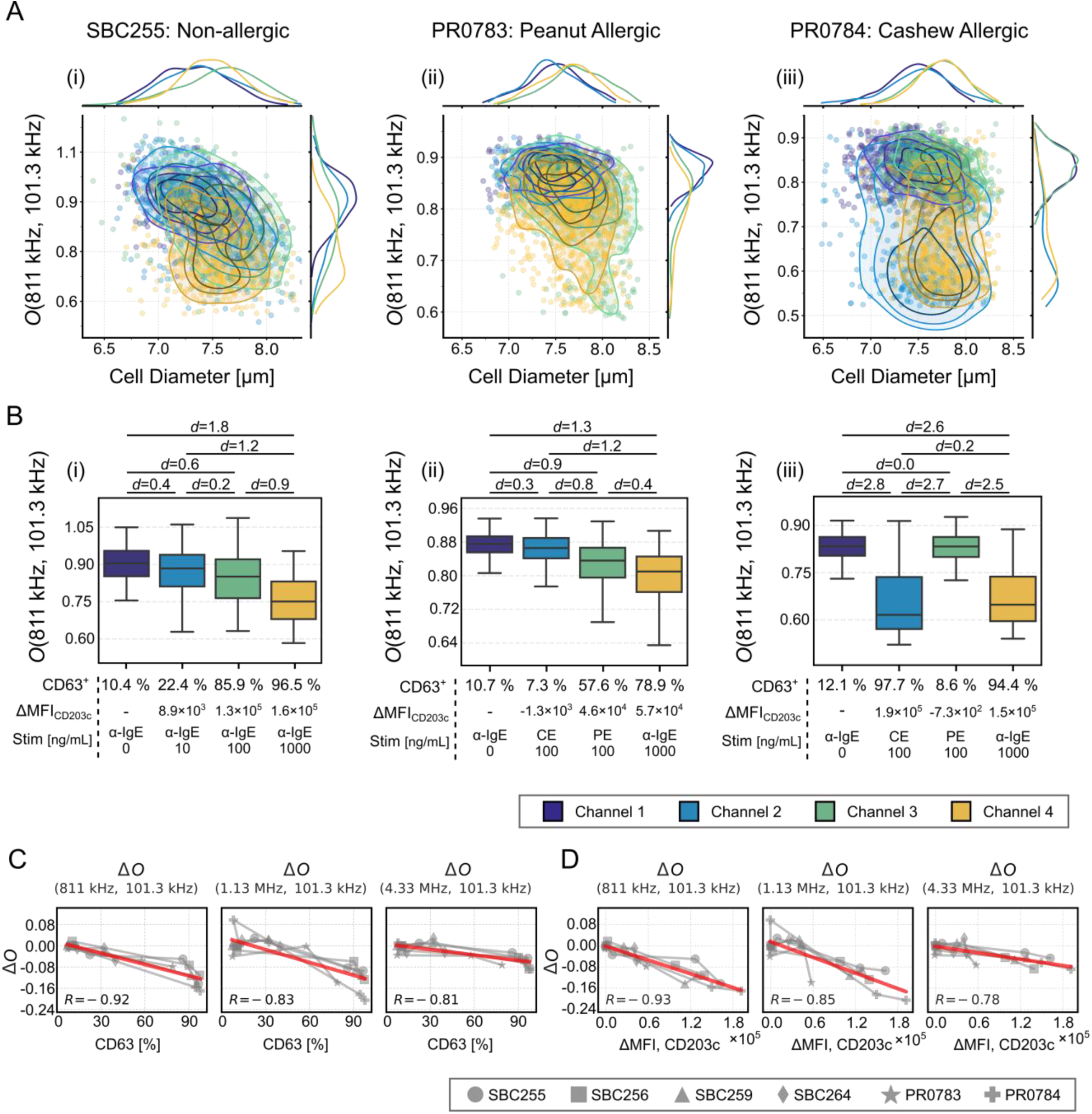
Label-free basophil activation measurement results. (A) Scatter plots of opacity 𝑂(811 𝑘𝐻𝑧, 101.3 𝑘𝐻𝑧) versus diameter from representative IFC measurements from three donors: (i) SBC255, (ii) PR0783, (iii) PR0784. The contour lines denote the 25^th^, 50^th^, 75^th^ percentiles of each population group, from the darkest to lightest shade. (B) Opacity 𝑂(811 𝑘𝐻𝑧, 101.3 𝑘𝐻𝑧) versus activation levels (%CD63^+^, and ΔMFI_CD203c_) determined by fluorescence flow cytometry for the same donors. Effect sizes (Cohen’s *d*) are reported for pairwise comparisons. (C)-(D) Opacity and size changes measured by impedance flow cytometry (IFC) compared with basophil activation levels measured by fluorescence flow cytometry. Among the six donors, four (SBC) were anonymous with unknown allergy status, and two were patients with allergy status confirmed by skin prick testing. Gray lines indicate changes in opacity or diameter for each blood sample, and red lines show the best-fit linear models for the corresponding data. Pearson correlation coefficient (R) values are indicated in the bottom left corner of each plot.

## 4. Conclusions

In summary, we developed a pipeline based on a deep unfolding network for processing code-multiplexed, multi-frequency IFC data. Our pipeline resolved singlet, doublet, and triplet overlaps in a cell-bead mixture dataset accurately. We inferred multi-frequency impedance profiles using least-squares fitting and achieved accurate reconstruction of impedance-intensity distributions across all measured frequencies. Using our pipeline, label-free physical profiling of basophil activation reproduced the activation-dependent opacity trend consistent with our previous study. Unlike prior approaches, our pipeline explicitly models bit-level intensity scaling and temporal stretching to handle nonlinearity in measured impedance signals. Furthermore, the unified network architecture improves generalization by sharing representations across tasks and simplifies scaling and maintenance of the network. To the best of our knowledge, this is the first demonstration of applying code multiplexing to multi-frequency IFC.

Although we demonstrated the initial technical feasibility of our method, this study has several limitations. First, the flow rate and the corresponding throughput of our current system was relatively low. The maximum flow rate was constrained by the dynamic response of the electrical measurement system and was kept low enough for the electronics to capture the rapid transit of cells through the constriction channel without degrading signal integrity. To overcome this challenge, it should be possible to use an oil-phase virtual constriction that preserves the electrical field constriction while avoiding rapid cell acceleration in the constriction region (Zhu et al., 2023). Second, the performance of the deep unfolding network was evaluated only with synthetic overlap data. Decomposing experimentally-acquired overlapping waveforms into their constituent signals requires an alternative way to verify the overlaps, and should in principle be possible with wide-field high speed imaging synchronized with the electrical measurements. Third, linear fitting used for multi-frequency inference may not remain valid for IFCs at high particle electrical volume fractions. The Maxwell mixture theorem, which underlies the linear fitting, is valid only for spherical particles at low electrical volume fractions (< 10%). At higher particle volume fractions, the permittivity of the particle-medium mixture couples to the particle volume fraction as descried in Bottcher-Polder-van Santen and Bruggeman-Hanai formulations. These formulations indicate that cross-frequency impedance profiles can deviate from linearity for particles at higher volume fractions (Raicu and Feldman, 2015). Accordingly, in regimes where a simple linear approximation no longer holds, additional analytical methods will likely be required. Fourth, the current study was restricted to a small number of anonymous healthy donors and allergic patients. To establish diagnostic relevance for allergy assessment, a larger cohort of allergic patients is needed.

Although further investigations are needed, our study demonstrates the technical feasibility of a pipeline based on a deep unfolding network and its potential to expand code-multiplexed IFC applications. While this work used length-4 codes, which ideally support 16-channel multiplexing, the principles of our pipeline apply to codes of other lengths as well. However, increasing code length or the number of channels lengthens the electrodes, raises their resistance, and increases sensor variation. Therefore, the practical limit should be determined empirically. Although we focused on the measurement of basophil activation for food allergy assessment, our pipeline could also facilitate IFC parallelization across diverse cell types and physiological readouts. For instance, given its ability to handle experimental nonlinearity, our pipeline may apply to complex IFC setups such as deformation cytometry. With further validation and optimization, our pipeline could enable parallelization of IFC-based multimodal single-cell measurements that combine electrical and mechanical properties.

## CRediT authorship contribution statement

Wonjun Lee: Writing – review & editing, Writing – original draft, Visualization, Validation, Software, Methodology, Investigation, Formal analysis, Data curation, Conceptualization. Sindy K.Y Tang: Writing – review & editing, Writing – original draft, Supervision, Resources, Project administration, Investigation, Conceptualization.

## Acknowledgement

This work was supported by the Department of Defense (DoD) Congressionally Directed Medical Research Programs (CDMRP) under Award Number: HT9425-23-1-1048. We thank Dr. Sungu Kim and Dr. Roslyn Massey for helpful discussions. The content is solely the responsibility of the authors and does not necessarily represent the official views of the Department of Defense.

## Data availability

Data will be made available on request.

